# RAMAN DEVELOPMENTAL MARKERS IN ROOT CELL WALLS ARE ASSOCIATED WITH LODGING TENDENCY IN TEF

**DOI:** 10.1101/2023.06.16.545357

**Authors:** Sabrina Diehn, Noa Kirby, Shiran Ben-Zeev, Muluken Demelie Alemu, Yehoshua Saranga, Rivka Elbaum

## Abstract

Tef (*Eragrostis tef* (Zucc.) Trotter) is an important staple crop in Ethiopia and Eritrea. Its grains are gluten-free and protein rich, so it is considered as a “super-food”. Adapting tef to modern farming practices could allow its intensive growth in other regions and enable larger communities to gain from its nutritional values. However, high lodging susceptibility prevents the application of mechanical harvest and causes significant yield losses. Lodging describes the displacement of roots (root lodging) or fracture of culms (stem lodging), forcing plants to bend or fall from their vertical position. Lodging is facilitated by various abiotic and biotic factors, and the lodging severity is increased in overpopulated fields. In this study, we aimed to understand the microstructural properties of crown roots, underlining tef tolerance/susceptibility to lodging. We analyzed plants at 5 and 10 weeks after emergence and compared trellised to lodged plants. Root cross sections from different tef genotypes were characterized by scanning electron microscopy, micro computed tomography and Raman micro spectroscopy. Lodging susceptible genotypes exhibited early tissue maturation, including developed aerenchyma, intensive lignification, and lignin with high levels of crosslinks. A comparison between trellised and lodged plants suggested that lodging itself does not affect the histology of root tissue. Furthermore, cell wall composition along plant maturation was typical to each of the tested genotypes independently of trellising. Our results suggest that it is possible to select lines that exhibit slow maturation of crown roots. Such lines are predicted to show reduction in lodging and facilitate mechanical harvest.

## 1. Introduction

With climate change and a steadily increasing world population, the improvement of cereal crops is a crucial task for plant breeders and farmers (Deutsch et al., 2018). Frequent events of unpredictable weather, such as higher temperatures and longer drought phases, demand urgent action to enhance crop production and resilience to stress conditions. Underutilized crops hold much potential for continued food production under unfavorable climatic conditions. A major cause of yield loss is lodging, defined as the permanent displacement of the plant from the erected position (Pinthus, 1974). This phenomenon is initiated by abiotic and biotic factors, like storms, soil texture and structure, high nitrogen levels causing excessive growth, or thin and weak stems that is more abundant in overpopulated fields. Lodging is divided into stem and root types (Berry et al., 2004). Stem lodging is the bending or breaking of the stem, while root lodging is the dis-anchoring of the plant from the soil, resulting in the stem falling without bending or breaking. Cultivation of dwarf and semi-dwarf plants is one of the most promising solutions for lodging (Berry et al., 2004). Studies on lodging are often focused on the major crops - wheat, maize, and barley (Berry et al., 2004, Sposaro et al., 2010, Berry and Spink, 2012, Piñera-Chavez et al., 2016).

Tef (*Eragrostis tef* (Zucc.) Trotter), is one of the most important staple crops in Ethiopia and Eritrea. Tef grains are gluten-free and protein-rich, considered a “super-food” (Girija et al., 2022), and its biomass is regarded as high-quality animal feed. Hence, this crop is gaining increasing interest worldwide. Lodging in tef is universal and poses a major cause of yield losses, particularly under mechanized and irrigated agricultural practices. Lodging can cause up to 50 % yield losses in addition to the reduction in the quality of grains (Paff and Asseng, 2018). In most cases, tef root lodging causes the displacement of plants, however, without uprooting the root system (van Delden et al., 2010, Ben-Zeev et al., 2023, Ben-Zeev et al., 2020). Similar to other crops, many studies are focusing on semi-dwarfism in tef(Bayable et al., 2020, Blösch et al., 2020). Nonetheless, recent studies indicate that rather than height, the root system architectures and individual roots mechanical properties may substantially contribute to lodging tolerance.

Cell walls, composed mainly of cellulose, hemicellulose, and lignin, determine the mechanical properties of plant tissues (Gierlinger and Schwanninger, 2007, Gierlinger, 2010, Simon et al., 2018, Gierlinger, 2018, Agarwal, 2006, Agarwal, 2010). Cellulose and hemicellulose are polysaccharides that exhibit limited composition variation. In contrast, the composition and structure of lignin, a polyphenolic polymer, varies among the plant species, organs, tissues, and even growth conditions (Liedtke et al., 2021, Zancajo et al., 2022). Lignin is deposited in mature cell walls to increase crosslinking between the cell wall polymers and stiffen the wall. Therefore, mapping lignin variation on tissue and cell levels is a remarkable target in understanding a plant’s biochemical composition. Raman micro-spectroscopy can provide fingerprint-like chemical information on a micrometer scale and thus form chemical images of plant microscopic cross sections. Recent spectroscopic studies about the lignin building blocks, such as coniferyl aldehyde, coumaric acid, ferulic acid, or dibenzodioxin, open up the possibility of mapping lignin variation within the polysaccharide cell wall via Raman micro-spectroscopy (Bock and Gierlinger, 2019, Bock et al., 2020).

In order to separate Raman from fluorescence signals and to interpret overlapping Raman bands, additional tools, such as background correction, Gaussian fitting, and normalization, are often applied to decompose the signals and assign bands to certain molecules or functional groups (Zancajo et al., 2022, Liu et al., 2015). Raman micro imaging of plant tissues present high heterogeneity on varied scales, indicating histological, and subcellular features (Agarwal, 2006, Zancajo et al., 2022, Piqueras et al., 2012, Baranska et al., 2006). Furthermore, analysis that includes many Raman maps from several samples increases the variation in a dataset. A sufficient data management and pre-processing is required to access variation that would carry biological meaning (Piqueras et al., 2014).

Our previous studies identified tef genotypes that are lodging tolerant while others that are lodging susceptible, possibly due to root-related traits (Ben-Zeev et al., 2023). Therefore, here we aimed to test (1) the association between lodging tendency and the composition of the cell walls in tef crown roots, and (2) whether the lodged position influences root anatomy, tissue and cell wall composition. Using bright-field light microscopy, micro computed tomography (μCT), scanning electron microscopy (SEM), Raman micro-spectroscopy, and univariate and multivariate analyses, we identified tissue and cell wall markers that develop with root maturation and plant lodging. Our results suggest that the lodging might be associated with early development of aerenchyma. In contrast, cell wall molecular makeup was found to reflect a genetic tendency to lodging independent of the plant position.

## 2. Materials and methods

### 2.1 Plant material and sample preparation

Tef plants of four genotypes (RTC-157, RTC-275, RTC-392, and RTC-400) with different tendencies for lodging in the field (Table S1) were sown in 3.9 l pots filled with professional potting soil (Tuff Golan, Merom Golan, Israel), grown in a greenhouse at 28/22 °C day/night and watered with tap water. After emergence, seedlings were thinned to one per pot, resulting in 72 tef plants in total. One third of the plants were harvested 5 weeks after emergence (5w), then a second third was trellised up with bamboo to prevent lodging (10wt) and the remaining third was growing free (10wf). The trellised and free-growing plants were harvested 10 weeks after emergence, when lodging occurs universally independent of the genotype (see Table S1). Random plants from the four genotypes were selected for further investigations. Figure S1 shows a set of plants analyzed, typical in this study. All 10wf plants lodged at harvest. Since lodging was shown to be correlated with flowering time (Ben-Zeev et al., 2023), the genotypes RTC-275 and RTC-400 that show similar flowering time (Table S1) were chosen for detailed comparison.

Crown and roots were cut into ca. 2 cm in length, rinsed with double distilled water, and fixed in formalin (50 % Ethanol, 2 % Acetic acid, 10 % Formaldehyde). Samples were stored at 4 °C until processed. Cross sections were taken from the proximal part of the crown roots (close to the crown) using a razor blade.

### 2.2 Scanning electron microscopy

The cross sections were placed on a metal stub covered by carbon tape. Images were obtained using Phenom (Thermo Fisher) SEM equipped with a secondary electron detector.

### 2.3 Micro tomography

A SkyScan 1272 system (Bruker microCT, Kontich, Belgium) was used for the X-ray micro-CT measurements. The crown roots were dehydrated in a graded ethanol solution series (70 %, 90 % for 1h, 100 % 1h) and then moved to 1 % w/w iodine in 100 % ethanol, and stored at 4 °C for at least 48 hours before the measurement. The roots were rinsed in ethanol to remove residuals of iodine and then loaded into plastic tube sample holders and mounted in the µCT-system. For each root, 1800 2D projection images were collected at 40 KV over 360° at two magnifications, resulting in a 6 μm^3^ and 7.99 μm^3^ voxel resolution. The raw tomographic images were reconstructed with the NRECON® software (Bruker, Belgium) and visualized using Fiji software (Schindelin et al., 2012)

### 2.4. Raman measurements

Cross sections were placed on a glass slide with a droplet of water, covered with a cover slip, and sealed with nail polish. Raman maps were acquired using InVia Raman microscope (Renishaw, New Mills, 178 UK) operated with WiRE3.4 (Renishaw), equipped with polarized 532 nm laser (45 mW), and a 63x water immersion objective. The cross sections were scanned using 1 μm steps in x and y direction. Spectra were collected with an 1800 lines/mm grating, resulting in a spectral resolution of ∼4 cm^-1^, and an integration time for 0.01 s each spectrum.

### 2.5. Data analysis

Data management, pre-processing, and analysis were executed in Matlab R2020b (The Mathworks, Inc., Natick, MA, USA). Data pre-processing included interpolation of the spectra using a distance of 1.8 cm^-1^ in the range from 350-1800 cm^-1^, followed by an asymmetric least-square background correction as proposed by Eilers (Eilers, 2003). Subsequently, the spectra were vector-normalized. For data analysis, Raman spectra from the cell walls were selected and cell lumen spectra were excluded from further investigation. To achieve the selection of cell wall spectra, the integrals of the area between 1550 and 1700 cm^-1^ in all preprocessed Raman spectra from each map were compared with an appropriate threshold. The bands in the investigated area are most common for lignin in the cell wall and therefore suitable for this selection (Liedtke et al., 2021).

Chemical mapping as well as multivariate analyses and imaging were applied on a dataset containing the selected pre-processed cell wall spectra from maps representing the root cross sections of individual plants. Hierarchical cluster analysis (HCA) was carried out using Euclidean distances and *Ward’s* algorithm (Ward, 1963). The images were created using assignments to the four biggest clusters of the dataset. Hypertools3 (Amigo et al., 2015) was used to create intensity maps of specific Raman shift, multiset principal component analysis (PCA), and multivariate curve resolution (MCR) (de Juan et al., 2014).

## 3. Results and discussion

### 3.1 Histological variations between genotypes along plant development

In the present study, we investigated crown roots of four tef genotypes at five weeks after emergence (5w) and trellised (10wt) and free (10wf) plants at 10 weeks after emergence (Figure S1). We compared roots of genotypes with low (RTC-157), intermediate (RTC-275, RTC-392), and high (RTC-400) tendencies to lodge (Table S1). Histological variations at the two development stages were examined by SEM (Figure 1). Roots of 5w plants exhibited no systematic variations in the thickness of cell walls, as expected for a young developmental stage. In contrast, the roots of 10-week-old plants showed a gradient of cell wall thickening from the center of the stele to the endodermis. In general, the stiffness of plant tissues increases with maturation, rendering mature roots stiffer and more fragile than young ones (Brule et al., 2016, Wang et al., 2012, Lichtenegger et al., 1999). Therefore, we expected the young roots to be flexible, and the older roots to fracture. Indeed, 5w tissues tended to collapse under vacuum in the SEM, possibly due to their high elasticity. The tissues of both trellised and free 10-week-old plants tended to fracture during sectioning (Figure 1, E-L).

**Figure 1:**
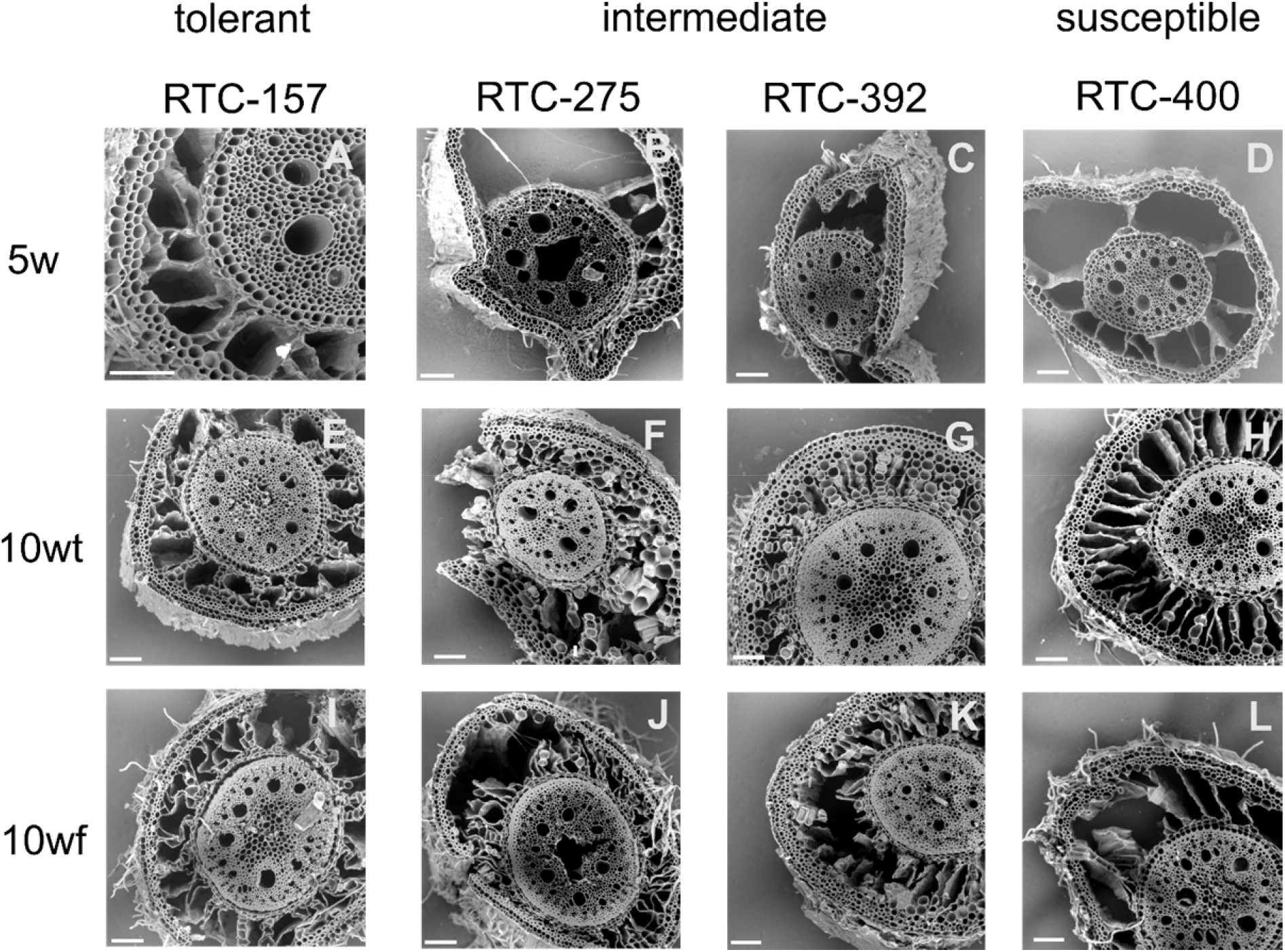
Representative scanning electron micrographs (SEM) of crown root sections taken from plants with varied genetic tendency to lodge, as indicated by the titles of the columns. **A-D**, Root sections of 5-week-old plants (5w). **E-H**, Root sections of 10 weeks old plants that were trellised to a bamboo (10wt). **I-L**, Root sections of 10 weeks old plants that were free to lodge (10wf). Scale bar, 100 μm.

Root cortex developed big holes termed aerenchyma. Markedly, aerenchyma was detected already at 5 weeks in the lodging susceptible genotype (RTC-400) (Fig 1D). Since aerenchyma may develop with plant maturation and transition to flowering, we aimed to compare RTC-400 to a genotype with similar phenology but more tolerant to lodging (Ben-Zeev et al., 2023). Our previous observations indicated that RTC-275 has flowering time similar to RTC-400 but intermediate lodging susceptibility (Table S1). Bright field images of hydrated cross sections demonstrated intact cortex with no development of aerenchyma voids in RTC-275 (Figure S2). To examine further the development of aerenchyma, we imaged roots via micro-computed tomography (μCT) (Figure 2). Similarly, μCT images showed aerenchyma development already at 5w samples in RTC-400, and delayed aerenchyma development at 10wf and 10wt in RTC-275. Interestingly, the big aerenchyma voids developed similarly along the sampling region, circa 5 cm in length from the root base. We thus concluded that small sampling variation in the distance from the root base were not expected to show variation in aerenchyma and that our cross sections are representative for the root developmental stage.

**Figure 2:**
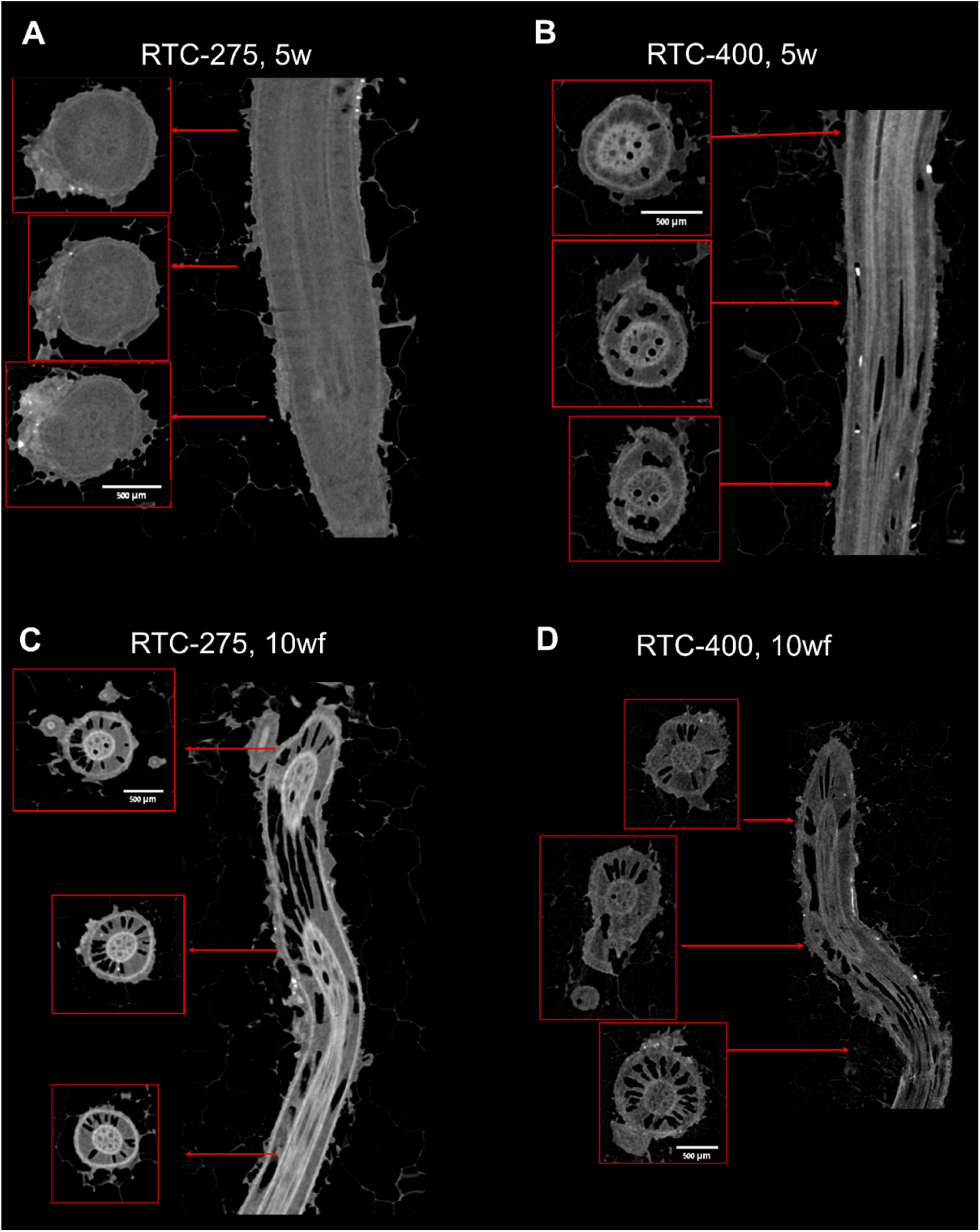
X-ray micro tomography (μCT) of crown roots. Roots of 5 weeks (5w) were imaged for RTC-275 (**A**), and RTC-400 (**B**). Roots of 10 weeks plants that were free to lodge (10wf) were imaged for RTC-275 (**C**) and RTC-400 (**D**). Scale bar, 500 µm.

In addition to aerenchyma, thickening of the inner tangential cell wall (ITCW) of the endodermis may also indicate root maturation (Fleck et al., 2011, Vaculik et al., 2009). The endodermis is a waxy barrier that prevents minerals and nutrients from diffusing freely between the soil solution and the plant sap. Its development starts with deposition of the lignified Casparian strips in the primary wall, followed by layers of secondary cell walls impregnated with suberin and lignin (Schreiber et al., 1999, Geldner, 2013, Song et al., 2019). SEM micrographs of the endodermis at 5 weeks show that the thickening of the endodermis did not start yet in both RTC-275 and RTC-400 (Figure 3A,B). In roots of 10-week-old plants, the endodermal ITCW thickened in both trellised and free plants (Figure 3C-F). However, the thickening was smaller in RTC-275 trellised plants (10wt) (Figure 3C). Collectively, our observations could not support histological variations between trellised and free plants. We suggest that while flowering time is similar between RTC-275 and RTC-400, the root aerenchyma may develop faster in RTC-400 than in RTC-275.

**Figure 3.**
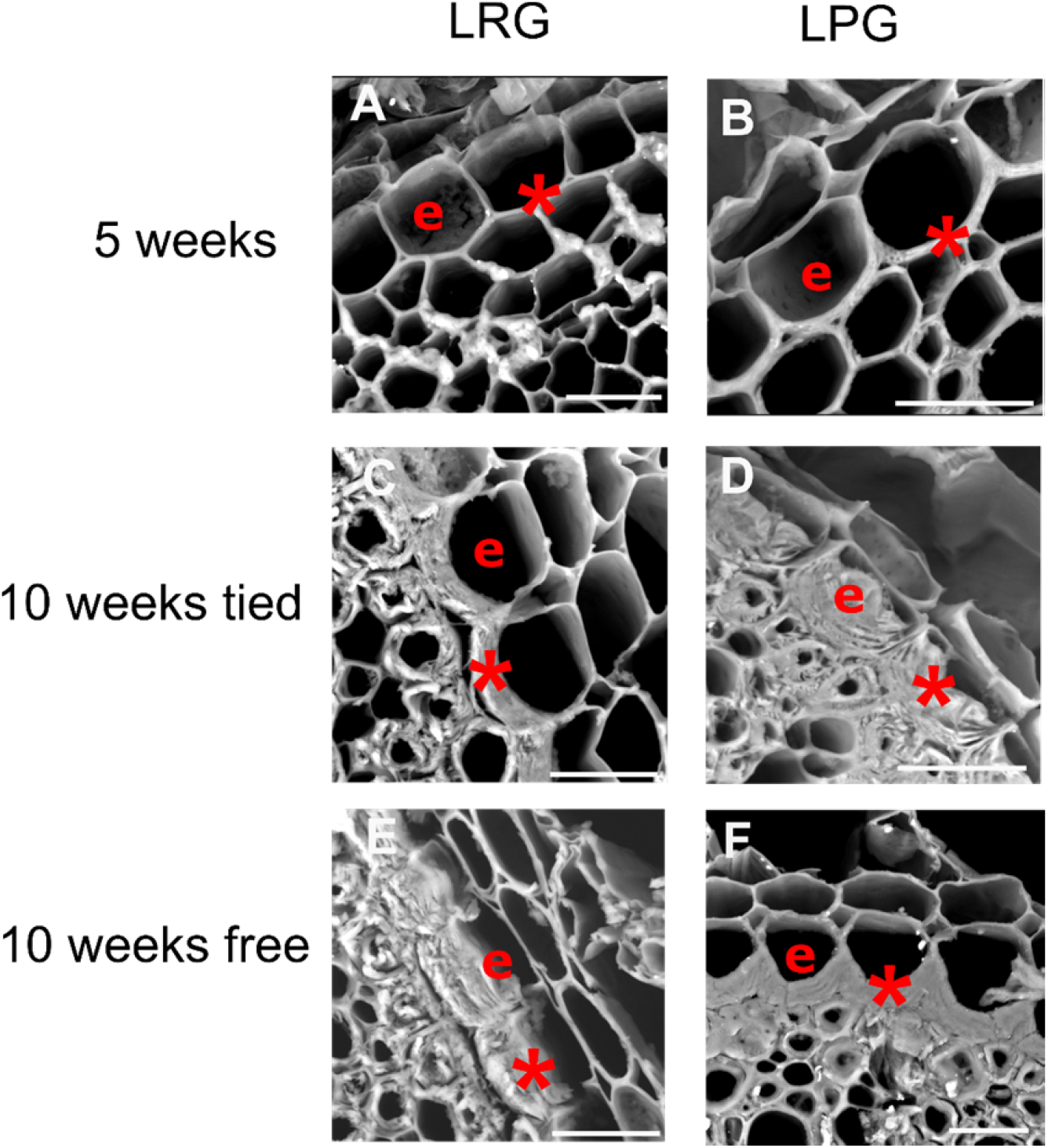
Representative scanning electron micrographs (SEM) of cross sections of the endodermis cell layer (e) of roots of 5 weeks (5w), 10 weeks trellised (10wt), and 10 weeks free (10wf) plants. A genotype with intermediate tendency to lodging (RTC-275, **A,C,E**) was compared to lodging susceptible genotype (RTC-400, **B,D,E**). The inner tangential cell wall is marked by asterisk (*). Scale bar, 20 μm.

### 3.2. Chemical composition of the root endodermis

#### 3.2.1. Mapping of specific Raman spectral bands

In order to monitor chemical variations within and between root sections, we applied Raman micro-spectroscopy. In the first stage, we collected Raman maps of the endodermis and stele of roots from RTC-400 and RTC-275 plants 10 weeks after emergence (Figure 4). Selected representative preprocessed spectra presented major spectral bands of lignin, namely, 1173 (coumaric acid/ ferulic acid), 1599 (phenolic lignin compounds), 1630 (lignin phenyl ring), and 1656 cm^-1^ (coumaric acid/ ferulic acid/ coniferyl alcohol) (Table 1). Cellulose and other polysaccharides contributed a band at 1084/1093 cm^-1^ (Agarwal and Ralph, 1997, Gierlinger et al., 2006). Bands assigned to suberin (at 1144 cm^-1^) (Heiner et al., 2018) and to asymmetric aryl ring stretch of lignin at 1510 cm^-1^ (Lupoi et al., 2015) were found only in the endodermis ITCW (Figure 4A). Bands at 1175 and 1209 cm^-1^, associated with coumaric and ferulic acid or H-units of lignin, were found only in xylem cell walls (Figure 4B). We also noted a blue shift in the lignin band at 1600 cm^-1^ in the xylem cell walls of 10wt roots. Table 1 summarizes the most abundant bands and their contributions in specific tissues. Raman intensity maps of selected peaks are presented in Figure 5. The cellulose-related band (1093 cm^-1^) showed little variation in its distribution between RTC-275 and RTC-400 under all growing conditions (Figure 5, top row). This suggested that the cell walls within each cross section are of comparable density. In contrast, the distribution of the suberin band at 1144 cm^-1^ indicated deposition in the endodermis inner tangential cell walls in all plants, except for the young RTC-275 plant (Figure 5, second row). Noticeably, in the RTC-400 10wf, suberin was detected also in the walls of the central cylinder external cell layer, termed pericycle (Figure 5Bf).

**Figure 4:**
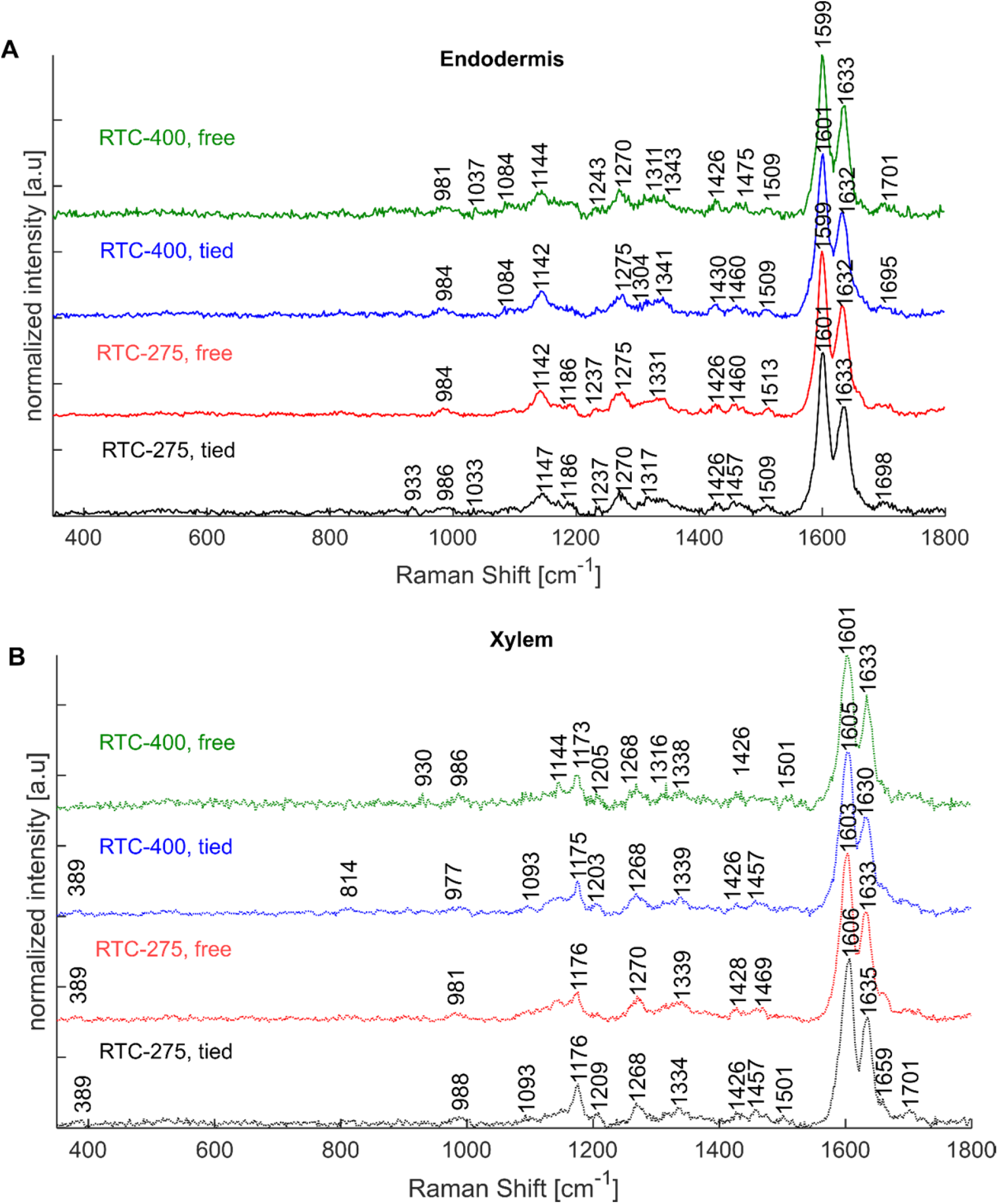
Representative pre-processed Raman spectra of crown root cross sections from 10-weeks-old plants of a genotype with intermediate lodging tendency (RTC-275) and a high lodging tendency (RTC-400). Spectra of the (**A**) endodermis and (**B**) xylem root tissues are compared. Colored arrows in Figure 5 indicate the location from which similar colored spectra were extracted.

**Figure 5:**
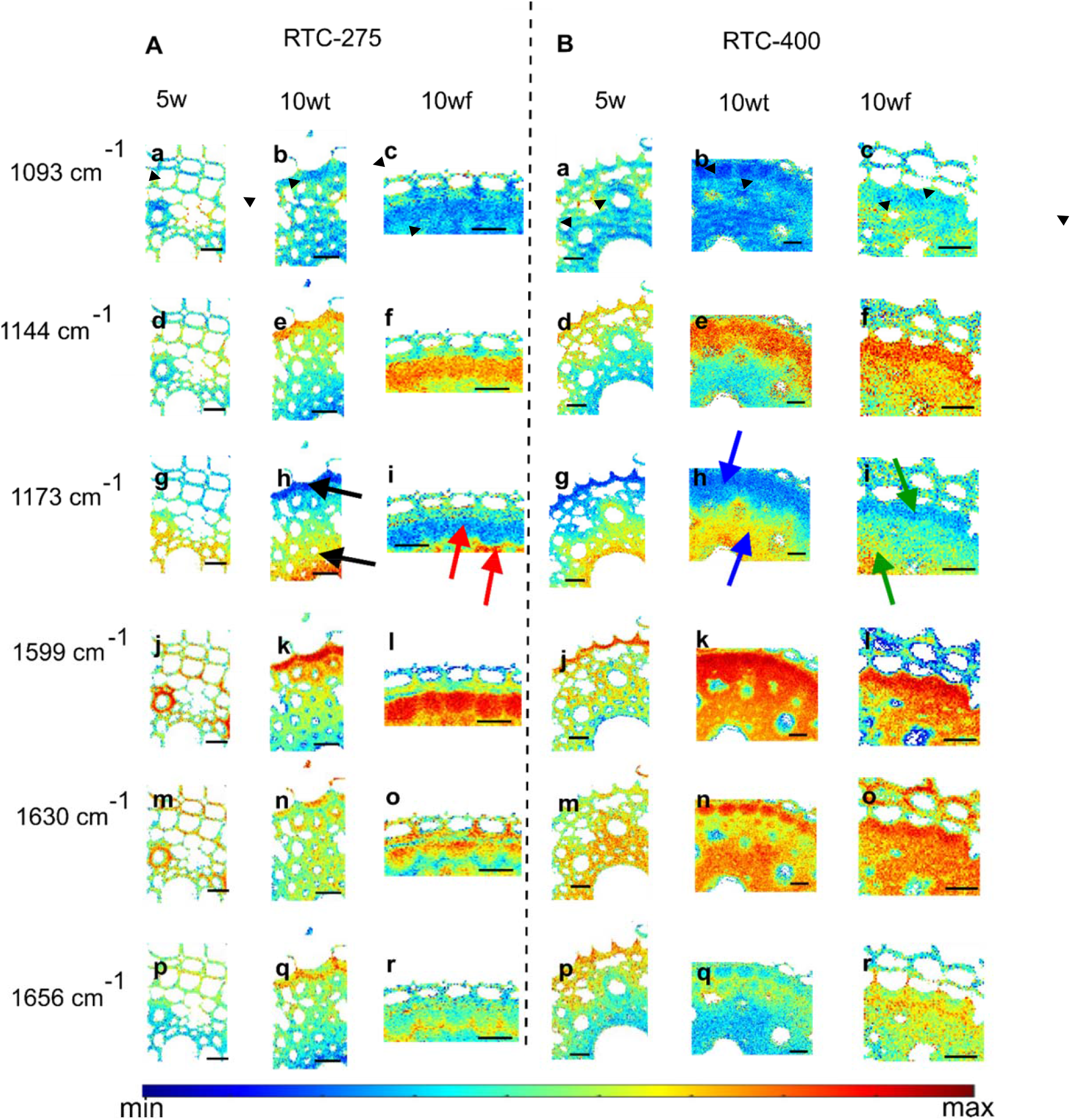
Intensity Raman maps collected at the endodermis and stele of crown root cross sections from a line with intermediate (**A,** RTC-275) and with high (**B,** RTC-400) lodging tendencies. The endodermis cell layer is marked in the top row by two arrowheads. Intensity maps are based on six Raman bands typical to major cell wall polymers: 1093 (cellulose, **a, b, c**); 1144 (suberin, **d, e, f**), 1173 (lignin, ferulic acid, coumaric acid, **g,h,i**); 1599 (phenolic ring, lignin **j,k,l**); 1630 (lignin, ferulic acid, coumaric acid, **m,n,o**); and 1656 cm^-1^ (coumaric acid, ferulic acid, coniferyl alcohol, **p,q,r**). The colored arrows represent the locations of spectra with similar colors shown in Figure 4. Scale bar, 20 µm.

**Table 1:** Raman bands in xylem and endodermis cell walls and their tentative assignments.

| Prominent bands<br>[cm<sup>-1</sup>] | tentative assignments | Abundant in tissue/maturity |
| --- | --- | --- |
| 1656 | coniferyl alcohol, coumaric acid, ferulic acid ($\nu(\text{C}=\text{C})$, $\nu(\text{C}=\text{O})$) (Bock and Gierlinger, 2019) | mature tissue,<br>first layer in endodermis |
| 1633 | lignin compounds (symm. stretching phenyl ring) (Agarwal, 2014, Hanninen et al., 2011)<br>coumaric acid (Sasani et al., 2021) | young tissue,<br>Stele and ITCW of endodermis |
| 1605-1610 | biphenyl lignin (aromatic ring mode) (Zancajo et al., 2022, Bock et al., 2020, Lupoi et al., 2015)<br>coumaric acid (Sasani et al., 2021) | fiber cell tissue |
| 1599 | lignin (symmetric ring stretch) (Agarwal, 2010) | mature tissue<br>endodermis |
| 1510 | lignin (aryl ring stretch, asymmetric) (Lupoi et al., 2015) | mature tissue<br>endodermis |
| 1457 | aliphatic compounds ($\delta(\text{CH}_2)$) (Yu et al., 2007) | |
| 1428 | lignin ($\delta(\text{O}-\text{CH}_3)$, $\gamma(\text{CH}_2)$) (Lupoi et al., 2015) | |
| 1339 | cellulose, lignin $\delta(\text{O}-\text{H})$ (Agarwal, 2019) | |
| 1270 | lignin (ring deformation) (Lupoi et al., 2015) |  |
| 1209 | lignin G-Units (C-C-H, C-O-H, aryl-O) (Yu et al., 2007), H-Units (Bock et al., 2020, Lupoi et al., 2015, Sun et al., 2011) | fiber cell tissue<br>xylem |
| 1186 | lignin biphenyl compounds $\delta(\text{C-H})$ (Bock et al., 2020) | |
| 1175 | coumaric acid, ferulic acid (Heiner et al., 2018, Sasani et al., 2021, Ma et al., 2014) | fiber cell tissue<br>xylem |
| 1144 | suberin (Heiner et al., 2018) | endodermis |
| 1125 | aliphatic compounds ($\nu(\text{C-C})$) (Yu et al., 2007) | |
| 1093 | cellulose/hemicellulose (Gierlinger et al., 2006, Agarwal et al., 2016) |  |
| 984 | cellulose/hemicellulose (Agarwal et al., 2011) |  |
| 930 | lignin (Bock et al., 2020, Lupoi et al., 2015) |  |
| 520 | cellulose/hemicellulose/lignin (Agarwal et al., 2016) |  |
| 384 | cellulose/hemicellulose (Agarwal et al., 2016) |  |

The 1173 cm^-1^ band, assigned to coumaric and ferulic acids, was dominant in the cylinder tissue with a decreasing intensity towards the endodermis, where this band vanished entirely (Figure 5, third row, in comparison to Figure 4). In contrast, the phenolic ring stretch band at 1599 cm^-1^ was particularly intensive in the endodermis ITCW (Figure 5, fourth row). An overlay of the two intensity maps revealed a negative correlation between 1599 and 1175 cm^-1^ bands, especially in RTC-275, 10wt (Figure S3A). Interestingly, the 1599 cm^-1^ band was positively correlated to suberin deposition in the endodermis, as indicated by the 1144 cm^-1^ band, suggesting that it may represent both lignin and aromatic regions in the suberin.

Negative correlation was found between the bands assigned to aromatic residues at 1630 and 1656 cm^-1^ (Figure 5, fifth and sixth rows). An early-deposited layer of the ITCW (at the cell wall periphery) was high in 1656 cm^-1^ band, while cell wall deposited later presented high intensity of the 1630 cm^-1^ band. This is enhanced when the two bands are overlaid, particularly in the RTC-275 10wt and RTC-275 10wf (Figure S3B). Note that this band is actually a small spectral shoulder (Figure 4). In conclusion, Raman spectroscopy enabled us to distinguish the endodermis based on specific spectral patterns and to map bands related to lignin monomers (1173 cm^-1^), suberin (1144, 1599 cm^-1^), and lignin (1599, 1630, 1656 cm^-1^). Importantly, our results indicated that genetic lodging tendency was correlated to early suberization of the root endodermis in tef plants, while trellising had no substantial effects on the cell wall composition.

#### 3.2.2. Principal component analysis (PCA) of 5w plants

To extend our insight regarding endodermis development in the context of lodging, we analyzed additional two genotypes exhibiting intermediate (RTC-392) and low (RTC-157) lodging tendency (Table S1). Figure 6 displays the results of a multiset principal component analysis (PCA) of the 5w RTC-275 and RTC-400 Raman maps presented in Figure 5 (first and fourth columns), together with Raman maps of root cross sections from RTC-157 and RTC-392 at 5w. Blue and red represent negative and positive score values for each principal component (PC), respectively. PC1 explained 27 % of the total variance and separated between the spectra based on a cellulose/lignin band at 1341 cm^-1^, and lignin bands at 1175, 1207, 1606, and 1632 cm^-1^, showing negative values (Figure 6Ba). Raman maps of the genotypes RTC-157 and RTC-275 showed mostly PC1 positive (red) values representing lower abundance of lignin bands, while RTC-392 and RTC-400 showed mostly negative (blue) values, indicating high impact of lignin (Figure 6Aa-d). This suggested a specific lignin composition that was more dominant in the cell walls of genotypes with high lodging tendency. Interestingly, in RTC-275 (Figure 6Ab) only one cell presented thick cell wall with negative (blue) PC1 score values. The abundant of this type of cells, possibly thick-walled fiber cell, was increased for the more lodging susceptible genotypes (Figure 6Ab-d).

**Figure 6:**
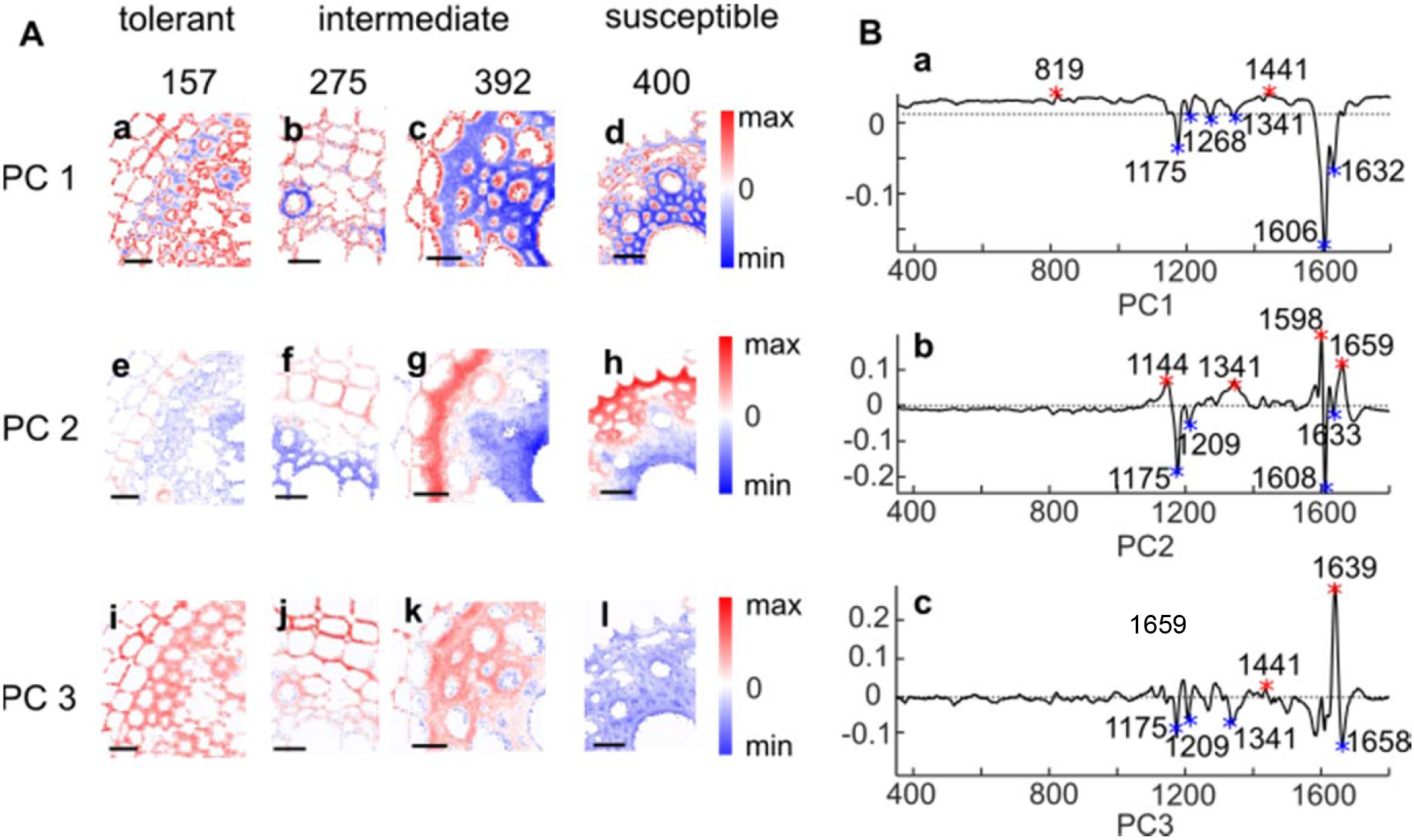
Principal component analysis (PCA) of Raman maps collected at the endodermis and stele of crown root sections extracted from plants of 5 weeks (5w). **A**, PCA images of root cross sections from four tef genotypes with varied lodging tendency. Subpanels in **A** represent distribution maps of PCs of (**a,e,i**) RTC-157; (**b,f,j**), RTC-275; (**c,g,k**) RTC-392, and (**d,h,l**) RTC-400. Images in **B** are the corresponding loadings presented for PC1 (**Ba**), PC2 (**Bb**), and PC3 (**Bc**). Blue pixels indicate negative PC score values and red pixel positive PC values. Scale bar, 20 µm.

PC2 explained 6 % of the total variance in the multiset, and represented an underlying pattern of Raman bands at 1144 cm^-1^ (suberin), 1341 cm^-1^ (cellulose/lignin), and 1598, 1659 cm^-1^ (lignin) (Figure 6Bb). These bands, and specifically suberin (1144 cm^-1^), are typical to developed endodermis cell walls, as discussed above (Figure 4, Table 2). Mapping score values of PC2 therefore highlighted the endodermis cell layer in red (Figure 6Ae-h). Noticeably, the intensity of positive PC2 values was increased with increasing lodging tendency of the genotype. The endodermis cell walls in RTC-157 and RTC-275 showed less intensive red pixels (Figure 6Ae,f). In the RTC-400 root, the pericycle was highlighted by PC2 mapping (Figure 6Ah). The steles of all genotypes presented mostly negative (blue) values of PC2. The corresponding loadings (Figure 6Bb) indicated that the lignin bands at 1175, 1209, and 1608 cm^-1^ were more intensive in the stele than the endodermis. While both stele and endodermis were lignified, we could identify a blue shift in the 1600 cm^-1^ band to 1608 cm^-1^, which is characteristic to the stele of the 5w plants. Similar shift was viewed in selected stele versus endodermis spectra of 10-week-old plants (Figure 4). Collectively, maps of PC1 and PC2 revealed that both lignification of the stele and suberization of the endodermis were increased in lodging susceptible relative to lodging tolerant lines.

Figure 6 Ai-l represents the PCA mapping results of PC3, which explained 3 % of the total variance (Figure 6Bc). The corresponding loadings revealed a strong influence of one band at 1639 cm^-1^, that could tentatively be assigned to lignin building blocks (coniferyl alcohol, ferulic acid, coumaric acid). The lodging prone RTC-400 showed only blue pixels (Figure 6Al). We assumed this band as a marker for lignin composition that was prevalent in younger tissue. Under this assumption, the RTC-400 showed an altered lignin composition associated with a more mature tissue already at 5 weeks.

### 3.3. Characterization of cell walls in the stele with respect to lodging tendency and plant maturity

To further establish the association between lodging and chemical maturation of cell walls, we added to our dataset older plants of 10wt and 10wf. We thus clustered the dataset through hierarchical cluster analysis (HCA) and mapped the four biggest clusters (Figure 7). Cluster 1 (Figure 7, black) was the largest cluster, having 10,079 spectra that were 44 % of the whole dataset; Cluster 2 (Figure 7, blue) contained 6,300 spectra that were 27 % of the dataset; Cluster 3 (Figure 7, red) contained 6,021 spectra that were 26 % of the dataset; and cluster 4 (Figure 7, green) contained only 635 spectra that were 3 % of the dataset. A dendrogram of those four clusters demonstrated that Cluster 1 and 3 were highly similar, and cluster 2 and 4 were more diverse (Figure 7B). The averaged spectra of each of the clusters presented strong lignin scatterings (Figure 7C). Noticeably, the averaged spectrum of cluster 1 presented a small lignin band at 1509 cm^-1^, which was stronger compared to the other average spectra. It further presented the suberin band at 1144 cm^-1^, missing from the other average spectra. In the averaged spectrum of Cluster 3, the bands at 1175 and 1207 cm^-1^ were more pronounced than in other average spectra (for band assignments see Table 2). Cluster 2 was typical to young 5w samples, with the exception of the RTC-392 5w map, in which the majority of spectra were associated with cluster 3. Spectra of cluster 1 were mostly distributed in maps of roots from 10wt and 10wf, with the exception of RTC-275 samples. These root sections are described by cluster 3. The smallest cluster 4 mapped majorly to 5w RTC-400. This cluster exhibited high variation and was mostly mapped to the cortex. We could thus identify the bands at 1509, 1209, and 1144 cm^-1^ in cluster 1 as typical to 10-week-old tissues that possibly represent maturation in the lignified fraction of the wall. In contrast, markers of young tissue were bands at 1605 and 1175 cm^-1^ in cluster 2 tentatively assigned to low degree of lignin polymerization, biphenolic compounds and hydroxycinnamic acids (Table 2). Noticeably, no systematic variation was identified between trellised and free 10-week-old plants, indicating that the cell wall composition was not influenced by the lodging itself, but was rather a genetically controlled trait. In order to identify possible chemical compositions that are responsible for the young versus mature Raman markers, we applied a common spectral unmixing approach to the same multiset. Multivariate curve resolution (MCR) (de Juan et al., 2014) with four components resulted in the distribution profiles and corresponding pure components presented in Figure 8. The first and second components could be assigned to cellulose (382, 988, 1096 cm^-1^) and young lignin (1175, 1207, 1270, 1339, 1458, 1605, 1633 cm^-1^). The signals at 1175, 1207, 1270, 1339, 1605, and 1633 cm^-1^ were associated with thick walled fiber cells (see discussion PCA, Figure 6, PC1). We termed the first MCR component as fiber-cell component I, which was abundant in 5w tissues of genotypes with increased lodging tendency, and in 10wt and 10wf tissues of RTC-157 and RTC-275 (Figure 8A,B). In general, fiber-cell component I was more intense in the inner (recently deposited) cell wall layers.

**Figure 7:**
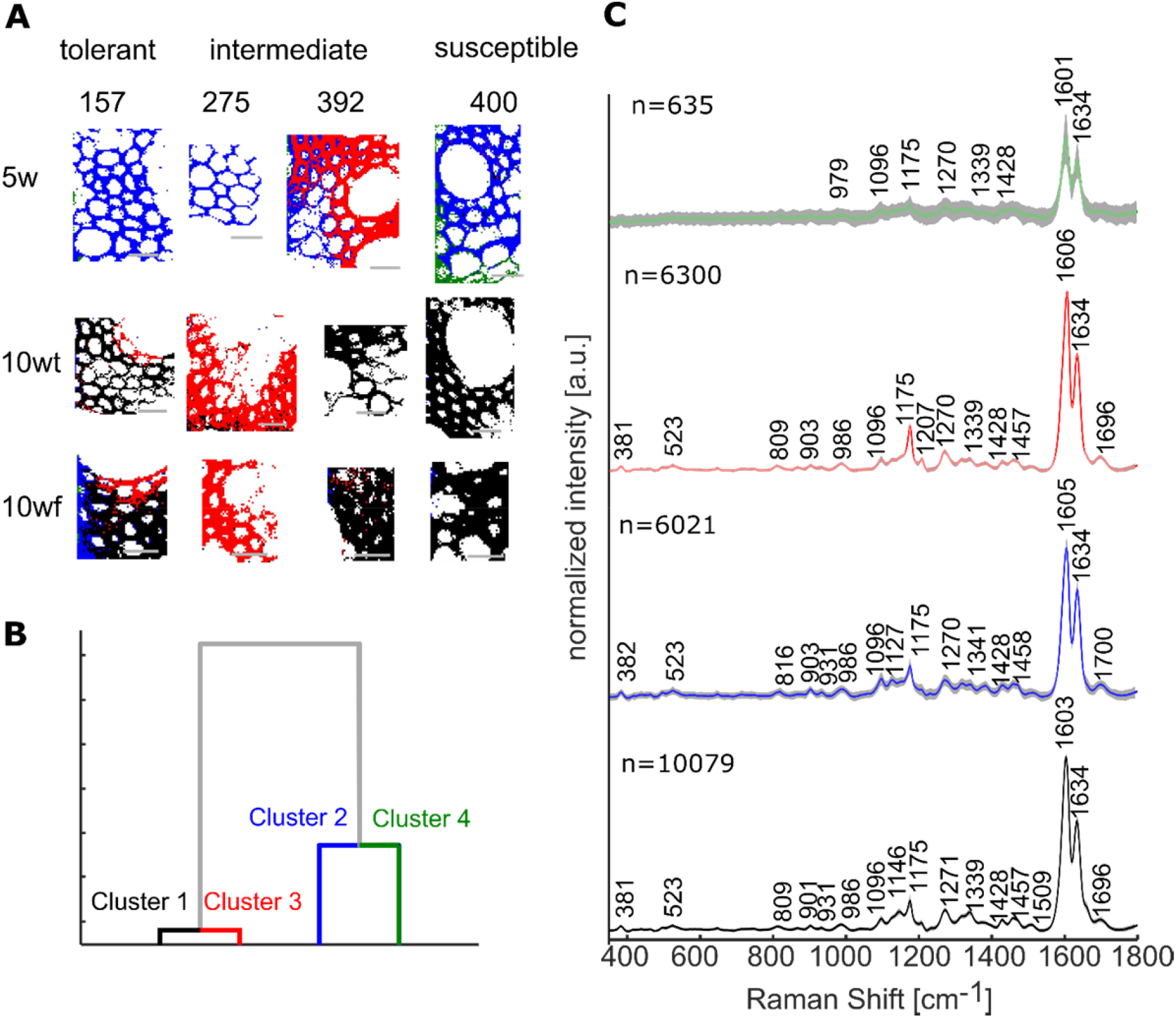
Hierarchical cluster analysis (HCA) of spectra from Raman maps collected from the stele of crown root cross sections of 5-week-old (5w), trellised 10-week-old (10wt), and non-trellised 10-week-old (10wf) tef plants of varied lodging tendencies. Based on Euclidean distance and *Ward’s* algorithm the spectra were grouped to four groups in which the variation within the groups is smaller than the variation between groups. **A**, Mapping of the four clusters. **B**, Dendrogram demonstrating the degree of similarity between the four clusters. **C**, Averaged spectrum of each of the clusters. The clusters are color-coded identically in all panels: cluster 1-black, cluster 2-blue, cluster 3-red, and cluster 4-green. Scale bar, 20 µm.

**Figure 8:**
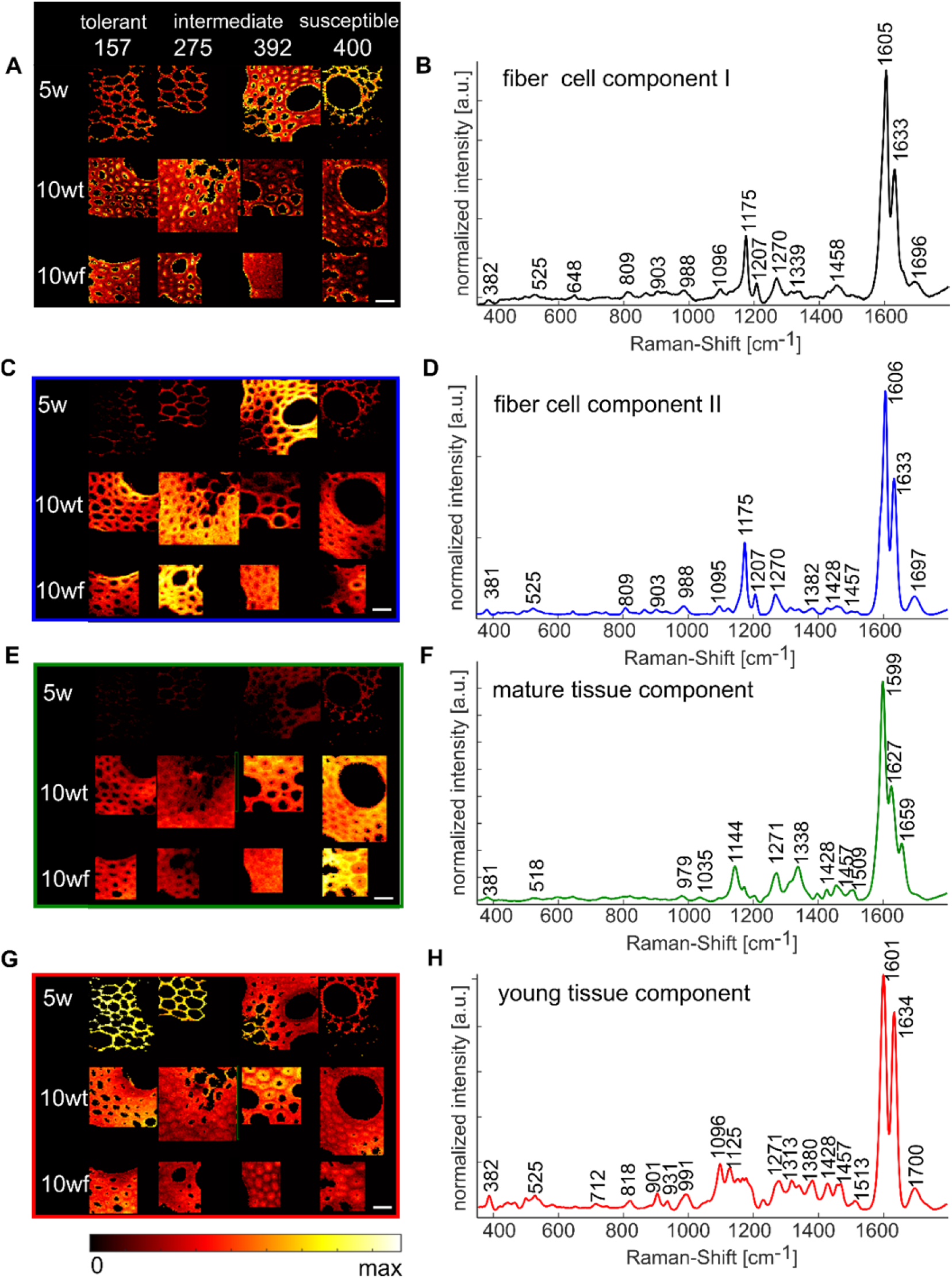
Spectral unmixing via multivariate curve resolution (MCR) of Raman mapping database of steles of crown root cross sections. Tef plants of 5-week-old (5w), trellised 10-week-old (10wt), and non-trellised 10-week-old (10wf) plants of varied lodging tendencies were examined. **A**, Heat maps of the pure component presented in **B. C**, Heat map of the pure component presented in **D. E**, Heat map of the pure component presented in **D**. **G**, Heat map of the pure component presented in **H**. Scale at the bottom of the figure represent relative intensity in all maps. Scale bars, 20 µm.

We termed the second component as fiber-cell component II, which showed only slight spectral differences in comparison to fiber-cell component I (Figure 8C,D): A small band at 1382 cm^-1^, which could be assigned to both cellulose and lignin, was more intense in fiber-cell component II than fiber-cell component I; A small shoulder at 1657 cm^-1^ on fiber-cell component I was missing in fiber-cell component II. Within specific cells that exhibited fiber-cell components, we noted that the distribution of fiber-cell component II was complementary to fiber-cell component I.

The third component was increasing with age and with increasing genetic lodging tendency (Figure 8E,F). We thus termed component 3 as mature-tissue component. This component presented lignin signals at 1035, 1144, 1271, 1338, 1428, 1457, 1509, 1599, 1627, and 1659 cm^-1^. In comparison to the fiber-cell components, the bands at 1144, 1338, 1509, 1627, and 1659 cm^-1^ were stronger in mature-tissue component. Furthermore, the main lignin band at 1605-1606 cm^-1^ in fiber-cell components was red-shifted to 1599 cm^-1^ in mature-tissue component, which may indicate higher degree of aromatic cross-linking in the mature-tissue versus fiber-cell components (Abraham et al., 2018). Interestingly, on a tissue level, the distribution of mature-tissue component was complementary to the distribution of the fiber-cell components, and appeared intensely in middle lamellas. We therefore suggest that the complex chemical structure termed ‘lignin’ changes along root maturation, and that this process is expressed in the spectral variations between the fiber-cell components and the mature-tissue component.

The fourth component was more intense in 5w samples, and negatively correlated to their lodging tendency (Figure 8H,G). We therefore termed it as young-tissue component. It included the lignin signals at 525, 931, 1271, 1380, 1428, 1457, 1513, 1601, 1634 cm^-1^ and cellulose at 382, 991, 1096, and 1313 cm^-1^, in addition to a band at 1125 cm^-1^, typical to lipids. The bands assigned to cellulose were stronger in this component compared to the fiber-cell components, possibly indicating low lignification levels. Interestingly, in this component, the intensity of the lignin band at 1634 cm^-1^ relative to the total lignin band at 1600 cm^-1^ was highest and calculated 0.85. In comparison, the same intensity ratio in both fiber-cell components was 0.6. The band at 1634 cm^-1^ could be assigned to lignin building blocks and indicate an initial stage of lignification in young tissues (see also Figure 6, PC 3).

## 4. Conclusion

Our analysis proposes a polymerization process of lignin during the stele maturation. In 5w samples, high contents of coumaric acid and other lignin monomers were indicated by the band at 1634 cm^-1^. This marked the increase in lignin building blocks in less lignified wall. These components polymerized during maturation of the wall, as was indicated in cross sections of 10-week-old plants by a relative decrease in the 1634 cm^-1^ band in parallel to the appearance of other lignin bands. The more mature tissue developed fiber cells and other cell walls with a specific lignin pattern, including bands at 1175, 1207, and 1605 cm^-1^. In contrast, highly mature tissues showed another specific lignin pattern, with bands at 1144, 1509, 1599, and 1659 cm^-1^. Here, lignification of the tissue is even more advanced, particularly causing a red shift of the total lignin band to 1599 cm^-1^ (Abraham et al., 2018). In this regard, roots of genotypes with high lodging tendency showed markers of mature lignin especially in the middle lamellae, while roots of more lodging-tolerant genotypes were less advanced in the lignification process and showed markers of fiber-cells components. We could not identify variation in cell wall maturation markers between supported and free plants.

In this work, we compared histological markers such as root aerenchyma and thickening of the endodermis ITCW and chemical composition of root cell walls between lodging tolerant and susceptible tef lines. Our histological analyses suggested that lodging might be typical to plants with advanced tissue development. In agreement with this, Raman micro-spectroscopy indicated early suberization and lignification of the endodermis cell walls of lodging susceptible genotypes. Furthermore, chemometric analyses revealed lignification of lodging susceptible genotypes and development of thick-walled fiber cells already at 5 weeks after seedling emergence. With the application of spectral unmixing tools, we identified three stages in lignin maturation, and could assign specific spectral features to each stage. Interestingly, we could not identify significant variations in the chemical composition of lignin between trellised plants and plants that were free to lodge, suggesting that the lodging itself did not influence lignin maturation. This would point to a genetic control of the process of lodging. It would be interesting to look for lignin maturation spectral signatures in other lignifying tissues and to mark developmental stages of lignification.

Overall, we claim that the roots of lodging-susceptible tef genotypes differed from those of more lodging tolerant ones. They developed aerenchyma, suberized endodermis, and lignified fiber cells at an earlier age, and the deposited lignin was cross-linked to a larger extent than plants of similar age that are more tolerant to lodging. High levels of aerenchyma, which makes the cortex fragile, together with lignified endodermis and stele, which make the central cylinder stiff, possibly challenge the ability of crown roots to resist plant lodging. Our results raise the hypothesis that stiff crown roots are more fragile and anchor the plants poorer than flexible crown roots. Further work may enable selection of tef genotype less susceptible to lodging.

## 5. Acknowledgements

S.D. is supported by a Minerva Fellowship of the Minerva Stiftung Gesellschaft für die Forschung mbH. S.B.-Z. is indebted to the Robert H. Smith Foundation. M.D.A. is The Heinrich Bonnenberg Scholarship Awardee. Y.S. is the incumbent of the Haim Gvati Chair in Agriculture. This research was partially supported by the ISRAEL SCIENCE FOUNDATION (grant No. 958/21).

## 6. Authors contributions

R.E., S.B-Z, S.D., and Y.S. designed the experiments. N.K controlled the growth experiment. M.D.A. and S.B.-Z. provided the background field data. N.K. and S.D. prepared the samples. S.D. acquired images using scanning electron microscopy and Raman spectroscopy. N.K. conducted the µCT measurements. S.D. performed univariate and multivariate analyses. R.E. and S.D. discussed the spectral data. All authors discussed the results. R.E. and S.D. wrote a first draft of the manuscript. N.K, S.B-Z, M.D.A and Y.S. reviewed and approved the manuscript. All authors agreed on the publication.

